# Platelet counts predict prognosis in IPF, but are not the main source of pulmonary TGFβ1

**DOI:** 10.1101/2020.03.06.978874

**Authors:** Deborah. L. W. Chong, Theresia A. Mikolasch, Jagdeep Sahota, Carine Rebeyrol, Helen S. Garthwaite, Helen L. Booth, Melissa Heightman, Ricardo J. José, Akif A. Khawaja, Myriam Labelle, Christopher J. Scotton, Joanna C. Porter

**Author notes:** These authors contributed equally (Christopher J. Scotton, Joanna C. Porter). Corresponding author: Dr Joanna C Porter Centre for Inflammation & Tissue Repair, Bloomsbury Campus, UCL, London, UK.

## Abstract

Transforming growth factor-β1 (TGFβ1) is the key pro-fibrotic cytokine implicated in the interstitial lung diseases (ILD), including idiopathic pulmonary fibrosis (IPF), but the primary source of TGFβ1 in these diseases is unknown. Platelets have abundant TGFβ1 stores, however their role in IPF is ill-defined. We sought to investigate whether platelets or platelet-derived TGFβ1 mediate IPF disease progression.

ILD/IPF and non-ILD patients were recruited to determine platelet reactivity and followed for mortality. To study whether platelet-derived TGFβ1 modulates pulmonary fibrosis, mice with a targeted deletion of TGFβ1 in megakaryocytes and platelets (TGFβ1^fl/fl^.PF4-Cre) were used in the bleomycin-induced PF model.

We found a significantly higher mortality in IPF patients with elevated platelet counts, along with significantly increased platelets, neutrophils, TGFβ1 and CCL5 in the lung and bronchoalveolar lavage (BAL) of ILD patients. Despite platelets being readily detected within the lungs of bleomycin-treated mice, neither the degree of pulmonary inflammation or fibrosis were significantly different between TGFβ1^fl/fl^.PF4-Cre and control mice.

Our results demonstrate for the first-time that platelet-derived TGFβ1 is redundant in driving pulmonary fibrosis in an animal model. However, platelets can predict mortality in IPF implicating other platelet-derived mediators, such as CCL5, in promoting human IPF disease.

## Introduction

The interstitial lung diseases (ILD) consists of inflammatory and fibrotic diseases affecting the lung interstitium [1], with idiopathic pulmonary fibrosis (IPF) representing the most severe of the idiopathic interstitial pneumonias (IIP) [2]. Transforming growth factor-β1 (TGFβ1) is a key mediator of fibrosis [3] and is elevated in bronchoalveolar lavage fluid (BALF) of IPF patients [4] and mouse pulmonary fibrosis (PF) models [5]. Genetic polymorphisms in *tgfb* are associated with IPF disease progression [6]. The source of TGFβ1 in IPF however is unknown, although alveolar macrophages, bronchial epithelial and PD-1^+^ TH17 cells have been proposed [7-9]. Identifying the exact cellular source is important as TGFβ1 is required for immunoregulation and tumour suppression and global deletion of this multi-functional cytokine would be detrimental.

Platelets are key mediators in immunity and tissue remodelling and represent an abundant source of TGFβ1 [10]. During pulmonary TB infection, platelets promote inflammation and extracellular matrix (ECM) degradation [11]. Platelets also attenuate liver fibrosis by degrading ECM [12], whilst platelet-derived TGFβ1 is pathogenic in cardiac fibrosis [13]. IPF patients have elevated mean blood platelet volume [14] and augmented platelet reactivity [15], whilst murine bleomycin-induced PF models exhibit increased platelet trapping in the lungs and collagen deposition [16]. Whether platelets are a major source of TGFβ1 in ILD and their role in mediating fibrosis is ill-defined.

Along with potential pro-fibrotic effects of platelet-derived TGFβ1, platelets may also stimulate neutrophil recruitment into the lung. Platelets and neutrophils act synergistically to cross inflamed endothelium [17]. Neutrophils and released mediators such as neutrophil elastase and neutrophil extracellular traps (NETs) have been linked to PF pathology [18-20]. It is unclear whether platelet-derived TGFβ1 impacts neutrophil recruitment into the lung *in vivo*.

The aim of this study was to define whether platelets and platelet-derived TGFβ1 mediate PF disease progression. We demonstrate that IPF patients with a higher blood platelet count (albeit still within the normal range), have a significantly higher mortality and we show increased numbers of platelets, neutrophils and TGFβ1 in the lung and BAL of ILD patients. However in the bleomycin-induced PF model using a conditional knockout transgenic mouse, where TGFβ1 has been deleted in megakaryocytes and platelets [21], platelet-derived TGFβ1 does not modulate disease severity. Interestingly CCL5; a platelet-derived chemokine, was significantly elevated in ILD patients’ blood and BAL, correlating with increased CXCL4 (platelet activation marker) and MMP7 (IPF disease severity biomarker). This suggests that other platelet-derived factors may promote human PF disease. To date, this is the first study where a specific cellular source of TGFβ1 has been addressed in the bleomycin-induced PF model and our findings build on, and refine, the current knowledge around platelets and platelet-derived TGFβ1 in mediating human PF disease.

## Materials and Methods

See supplementary materials for detailed methods

### Patient samples and mice

Human neutrophils, Platelet-rich plasma (PRP), BALF or lung biopsies from healthy controls, ILD or non-ILD patients were collected with written informed consent and research ethics committee approval. For the survival analysis, IPF patients (*n*=214) from the UCLH ILD Service were followed prospectively for mortality based on blood platelet counts as described in supplementary methods. TGFβ1^fl/fl^.PF4-Cre mice were obtained from Prof Richard Hynes [21] and were maintained with accordance to UK Home Office Regulations.

### Patient BALF & plasma sampling

BALF and plasma were collected and processed as described in supplementary methods. Cell counts were performed on cytospun BALF cells (*n*=6-12). Percentage of platelets in BALF (*n*=3-6) was quantified on a BD FACSVerse flow cytometer (BD Biosciences, San Jose, USA) and analysed with FlowJo v10 software (FlowJo LLC, Ashland, USA).

### Cytokine analysis

Total TGFβ1, PDGF-BB, CCL5, CXCL4 and MMP7 were quantified in PRP (*n*=4-5), plasma (*n*=11-47) or BALF (*n*=6-19) by ELISA (R&D systems, Minneapolis, USA). Active TGFβ1 was quantified by a Mink lung epithelial cell (MLEC) bioassay [22].

### Neutrophil chemotaxis assay

Human or murine neutrophils and PRP were isolated as described in supplementary methods. In brief, untreated or 1 μM SB-525334 ALK5 inhibitor (Sigma-Aldrich, Gillingham, UK) or DMSO pre-treated neutrophils (*n*=4) were added to the top of 8 μm PET transwells (Corning, New York, USA) and chemoattractants: 100 nM fMLP (Sigma-Aldrich), TGFβ1 (R&D systems), 10 ng/ml LTB_4_ (Cambridge Biosciences, Bar Hill, UK) or PRP added to the bottom. After 1 hour at 37°C, migrated cells were counted on a BD FACSVerse flow cytometer.

### Bleomycin animal model of pulmonary fibrosis

11-24 week old male and female TGFβ1^fl/fl^.PF4-Cre or littermate control mice (*n*=4-15) were given 25 or 50 IU bleomycin in a volume of 50 μl 0.9% saline, or saline alone, by oropharyngeal instillation. Mice were monitored for disease severity and culled at 6, 21 or 28 days post instillation.

### Quantification of inflammation in murine lungs

BALF was obtained from lungs following 3 washes of 1 ml PBS. BALF was centrifuged and cell pellet processed for cytospin cell counts. Murine lungs were homogenised in PBS before RBC lysis. Cells were blocked with Fc block (BD Biosciences) and stained with Ly6G-PE (clone 1A8), CD11b-APC (clone M1/70) and CD11c-APC-H7 (clone HL3) antibodies (all from BD Biosciences). Cells were analysed on a BD FACSVerse flow cytometer and FlowJo software.

### Quantification of fibrosis in murine lungs

Degree of fibrosis in lungs was assessed by micro-CT analysis [23]. Total lung collagen was quantified via the measurement of hydroxyproline by reverse phase HPLC [24].

### Histology & Immunohistochemistry (IHC)

Murine or human lungs were processed for histological analysis as described in supplementary methods. IHC was performed for murine (clone AB-7773, Sigma-Aldrich) or human CD61 (clone 2f2, Leica Biosystems, Wetzlar, Germany). Sections were scanned on a Nanozoomer Digital Slide Scanner and analysed using NDP.view software (both from Hamamatsu Corporation, Hamamatsu City, Japan)

### Statistical analysis

Survival analysis (multivariate cox proportional hazard regression, Kaplan-Meier curve and log rank test for trend) was performed using Stata v15.1 software (StataCorp LLC, Texas, USA). All other statistical analysis was performed with Prism v6 software (GraphPad, San Diego, USA) using linear regression, Mann Whitney U test, 1- or 2-way ANOVA with Holm-Sidak post-hoc testing. Mean ± SEM are shown in all graphs with a *p*-value <0.05 considered significant.

## Results

### Blood platelet counts predict mortality in IPF and TGFβ1 levels correlate with an activated blood platelet signature

We initially sought to investigate the role of platelets in IPF disease progression. A cohort of consecutive, unselected IPF patients (Supplementary table 1) were prospectively followed for mortality. They were split into three equally sized cohorts based on platelet count. Mortality was highest in patients with the highest platelet count (>267×10^9^/L), and lowest in those with the lowest platelet counts (<207×10^9^/L) (Log rank for trend *p*=0.04) (Fig 1a). Multivariate cox proportional hazard regression, correcting for age, sex and lung function (percentage of predicted Forced Vital Capacity) resulted in a hazard ratio of 1.25 (*p*=0.047) between platelet groups with the higher the platelet count, the higher the risk of mortality. This finding supports an association between platelets and IPF disease progression.

**Figure 1.**
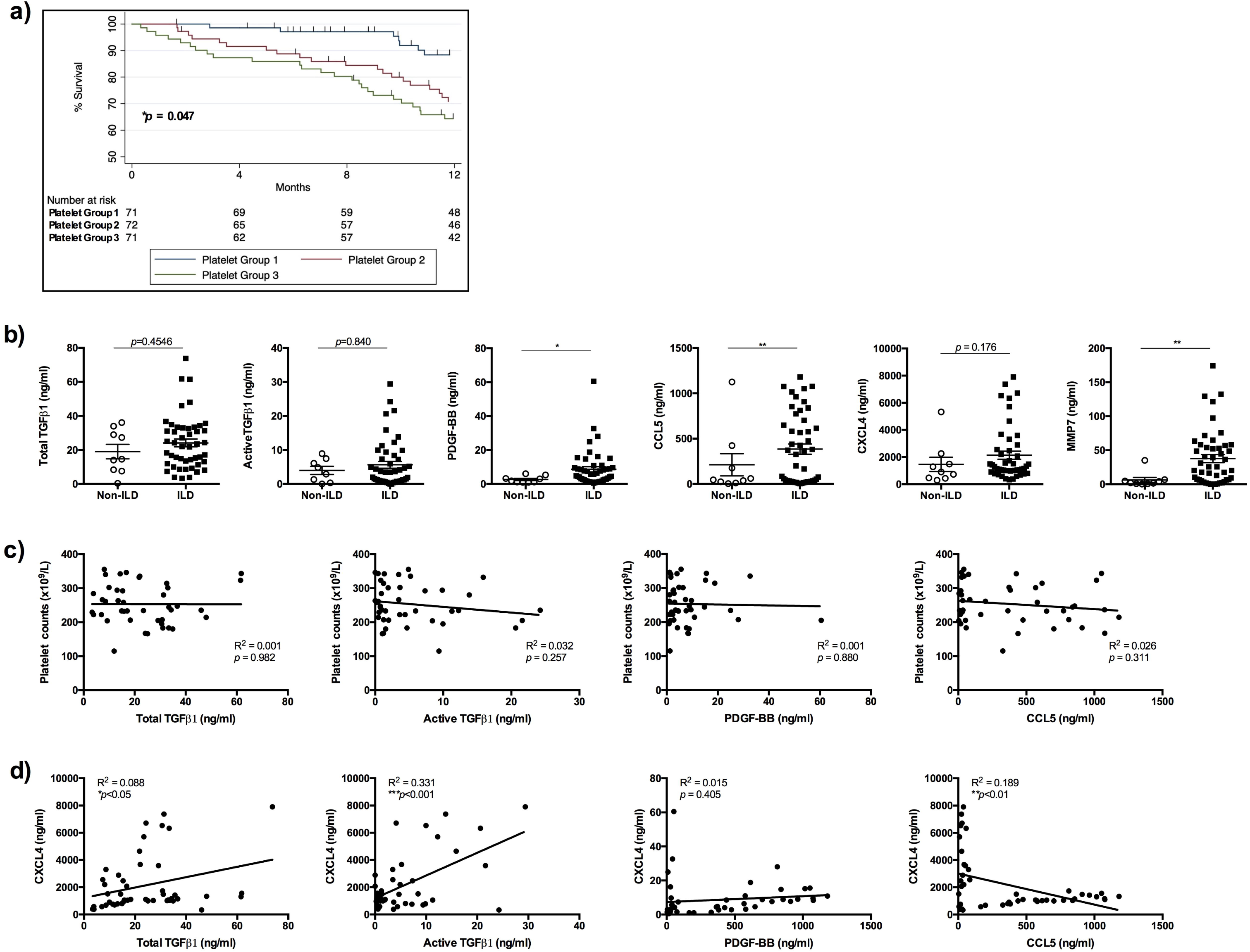
Human blood platelet counts predict mortality in IPF and levels of TGFβ1 correlate with an activated blood platelet signature. a) Kaplan-Meier survival curve of IPF patients (*n*=214) divided into three strata according to platelet count. The three strata were defined bias-free by dividing into three equal groups: Group 1=lowest third (blue), Group 2=middle third (red) and Group 3=highest third (green). Censored patients are indicated on the graph as ticks. Number of patients at risk for each time-point are shown below the Kaplan-Meier curve. b) Concentrations of total or active TGFβ1, PDGF-BB, CCL5, CXCL4 and MMP7 were measured in non-ILD and ILD patients’ plasma (*n*=11 and 47 respectively) by ELISA or MLEC bioassay. c) Concentration of mediators in ILD patients’ plasma were correlated with paired blood platelet counts (*n*=47). d) Concentration of mediators in ILD patients’ plasma were correlated with paired CXCL4 levels (*n*=47). Any statistical differences were determined using log rank test for trend, Mann Whitney-U test or linear regression and any significant differences are indicated (n/s = not significant, \**p*<0.05, \*\**p*<0.01, \*\*\**p*<0.001).

To investigate if platelets are a likely source of pro-fibrotic TGFβ1 in ILD, ILD and non-ILD patients were recruited (Supplementary table 2) to quantify platelet-derived mediators. Total and active TGFβ1 and CXCL4, a platelet activation marker [25] were increased in ILD plasma compared to non-ILD controls but did not reach significance (Fig. 1b). Platelet-derived growth factor-BB (PDGF-BB), CCL5 and MMP7, an IPF disease severity biomarker [26] were significantly elevated in ILD plasma (Fig. 1b). Total and active TGFβ1, PDGF-BB and CCL5 levels correlated poorly with paired ILD blood platelet counts (Fig. 1c) suggesting that high levels of these mediators were not due to increased platelet counts. However, total and active TGFβ1 in ILD plasma positively correlated with matched CXCL4 levels (Fig. 1d), whilst CXCL4 did not correlate with PPDGF-BB and negatively correlated with CCL5 (Fig. 1d). This suggests that activated platelets as represented by increased CXCL4 may be a source of TGFβ1 in ILD blood.

### Platelet-derived TGFβ1 is redundant in driving fibrosis or disease resolution in bleomycin-induced PF

Platelets are a source of TGFβ1 [10] with active TGFβ1 detected in thrombin-activated human PRP (Supplementary Fig. 1). To address whether platelet-derived TGFβ1 has a pro-fibrotic role, a conditional knockout transgenic mouse, in which TGFβ1 is deleted in megakaryocytes and platelets (TGFβ1^fl/fl^.PF4-Cre) [21] was used in the bleomycin-induced PF model. Unstimulated or thrombin-activated TGFβ1^fl/fl^.PF4-Cre PRP contained little or no active TGFβ1 compared to littermate PRP (Fig. 2a), confirming the knockout phenotype. 21 days post bleomycin treatment, littermate and TGFβ1^fl/fl^.PF4-Cre mice had regained initial weight loss with no difference between bleomycin-treated littermate or TGFβ1^fl/fl^.PF4-Cre mice (Fig. 2b). No difference was found between saline treated groups (Fig. 2b). Micro-CT analysis revealed fibrotic lesions within the bleomycin-treated lungs (Fig. 2c), with no significant difference in the percentage of fibrotic lung tissue or lung volume between littermate and TGFβ1^fl/fl^.PF4-Cre mice (Fig. 2c). HPLC analysis for hydroxyproline (a major component of collagen) content in whole lungs (Fig. 2d) confirmed that the loss of platelet-derived TGFβ1 did not impact fibrosis in this model.

**Figure 2.**
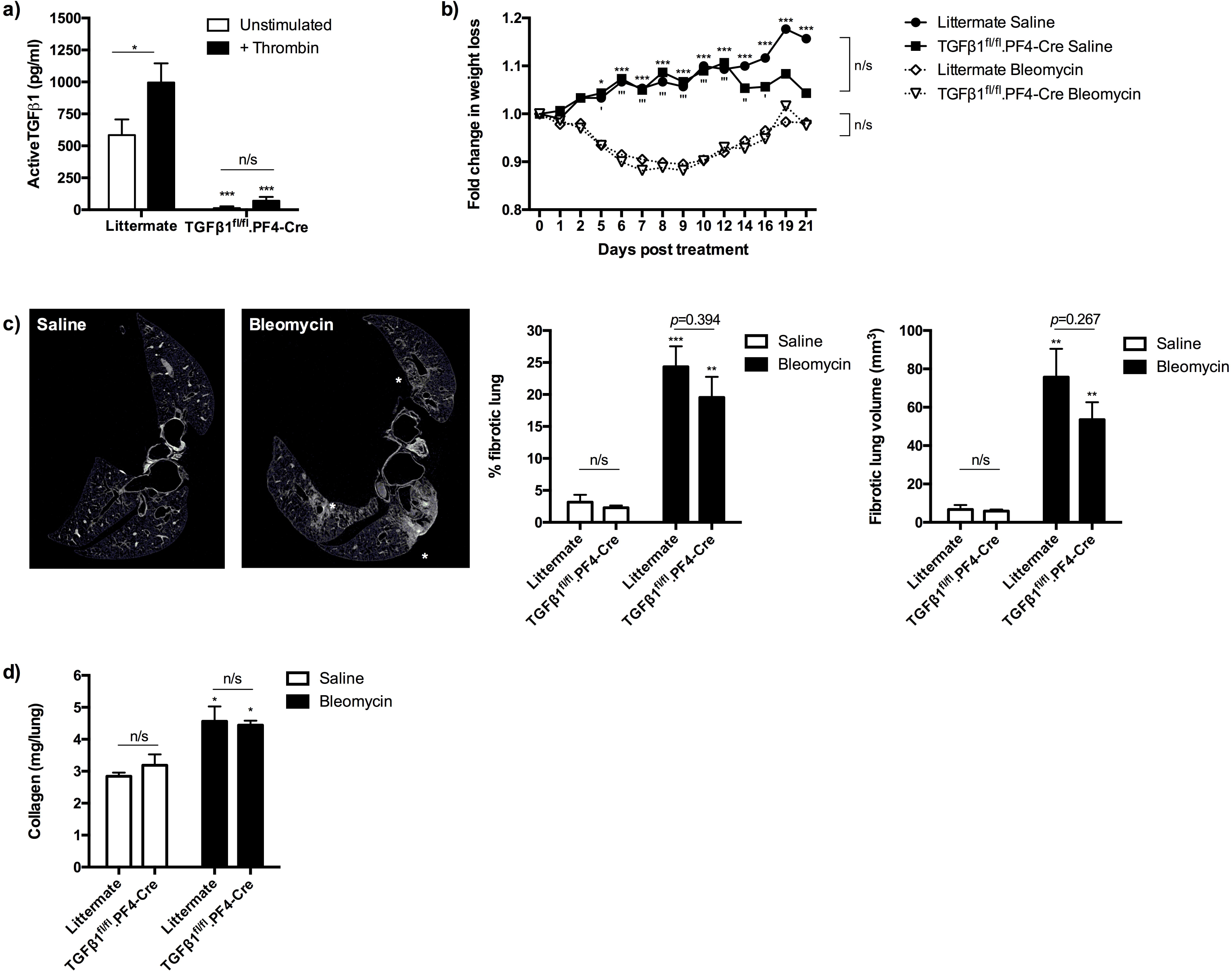
Platelet-derived TGFβ1 does not play a significant role in bleomycin-induced fibrosis. a) Quantification of active TGFβ1 by MLEC bioassay in unstimulated or thrombin-treated PRP from littermate (*n*=4) or TGFβ1^fl/fl^.PF4-Cre (*n*=5) mice. b) TGFβ1^fl/fl^.PF4-Cre or littermate mice were given 50 IU bleomycin (*n*=5/group) or saline (*n*=3/group) via an oropharyngeal route. Body weight was monitored for 21 days post instillation. c) Representative micro-CT scans of lungs after 21 days of treatment (white stars denote fibrosis). Percentage and lung volume of fibrosis of micro-CT scanned lungs was determined using InForm analysis software. d) Total lung collagen of murine lungs was determined by reverse-phase HPLC. Any statistical differences were determined using 2-way ANOVA with Holm-Sidak post-hoc testing or Mann-Whitney U tests. On Fig. 2b, asterisks represent significant differences between treatments in littermate control mice, apostrophes represent significant differences between treatments in TGFβ1^fl/fl^.PF4-Cre mice. On Fig. 2c-d, asterisks above the bars represent significance between saline and bleomycin treatment. Any other significant differences are indicated (n/s = not significant, \**p*<0.05, \*\**p*<0.01, \*\*\**p*<0.001).

Platelets contain two pools of latent TGFβ1; one released immediately upon activation and the latter slowly released during wound healing [27]. Administration of lower bleomycin doses allows the lung to heal after the initial injury. We examined whether platelet-derived TGFβ1 modulates disease resolution after bleomycin-induced PF. At 28 days after a lower dose of bleomycin challenge (25 IU), the percentage weight loss between littermate and TGFβ1^fl/fl^.PF4-Cre animals did not differ (Fig. 3a). Micro-CT analysis revealed equivalent remaining lung fibrosis (Fig. 3b-c). This suggested that platelet-derived TGFβ1 did not alter the rate of disease resolution. Histological analysis of lung tissue revealed dense cellular infiltration (as shown by H&E staining), ECM deposition (as shown by blue modified trichrome staining) and abundant CD61 expression (a platelet and platelet aggregates marker) in inflamed and fibrotic lesions (Supplementary Fig. 2). These results establish that platelet-derived TGFβ1 is redundant in driving lung fibrosis or disease resolution despite activated platelets represent an abundant source of pro-fibrotic TGFβ1 *ex vivo*.

**Figure 3.**
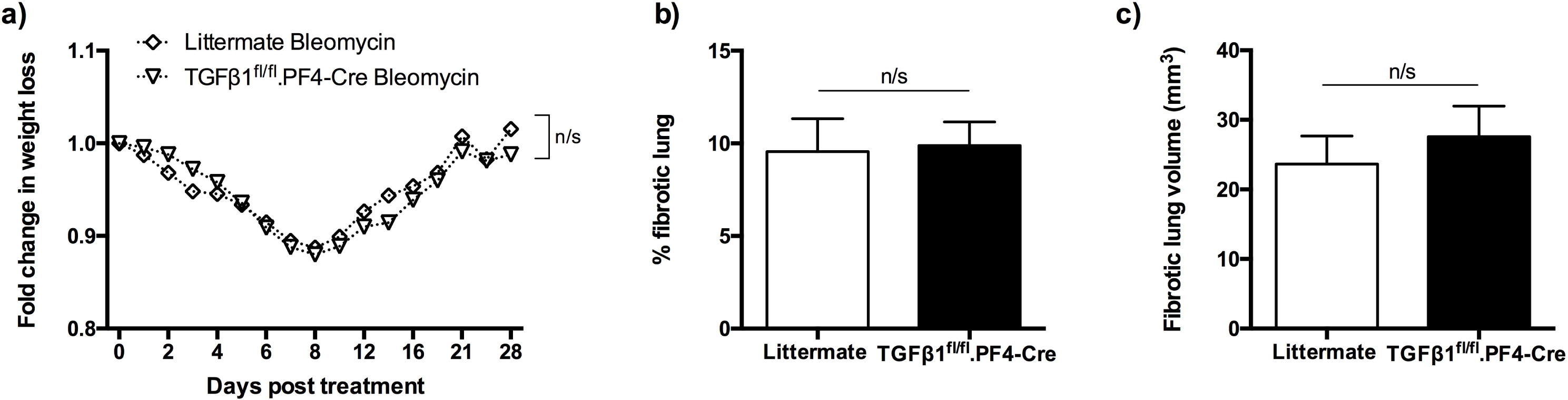
Platelet-derived TGFβ1 does not play a significant role in disease resolution after bleomycin-induced injury. a) TGFβ1^fl/fl^.PF4-Cre (*n*=8) or littermate mice (*n*=10) were given 25 IU bleomycin via an oropharyngeal route. Body weight was monitored for 28 days post instillation. b-c) Percentage and lung volume of fibrosis from micro-CT scanned lungs were determined using InForm analysis software. No statistical differences were found using 2-way ANOVA with Holm-Sidak post-hoc testing or Mann-Whitney U tests (n/s = not significant).

### TGFβ1 and platelet-derived TGFβ1 are potent neutrophil chemoattractants in vitro

Platelets aid neutrophil migration [17] but it is unknown whether platelets drive neutrophilic inflammation in ILD. Increased CD61^+^ platelets and platelet aggregates were found within IPF lung, particularly in the blood vessels (Fig. 4a). Interestingly, we also observed ILD BALF contained significantly more neutrophils than non-ILD controls (Fig. 4b). Due to increased neutrophil and platelet infiltration in ILD patients, we next assessed whether TGFβ1 influences neutrophil migration. Human neutrophils migrated in a concentration-dependent manner towards TGFβ1 (Fig. 4c) in an *in vitro* neutrophil chemotaxis assay. This effect was significantly reduced when TGFβ-receptor signalling was blocked by pre-incubating neutrophils with an ALK5 inhibitor (SB-525334) compared to DMSO vehicle controls (Fig. 4c).

**Figure 4.**
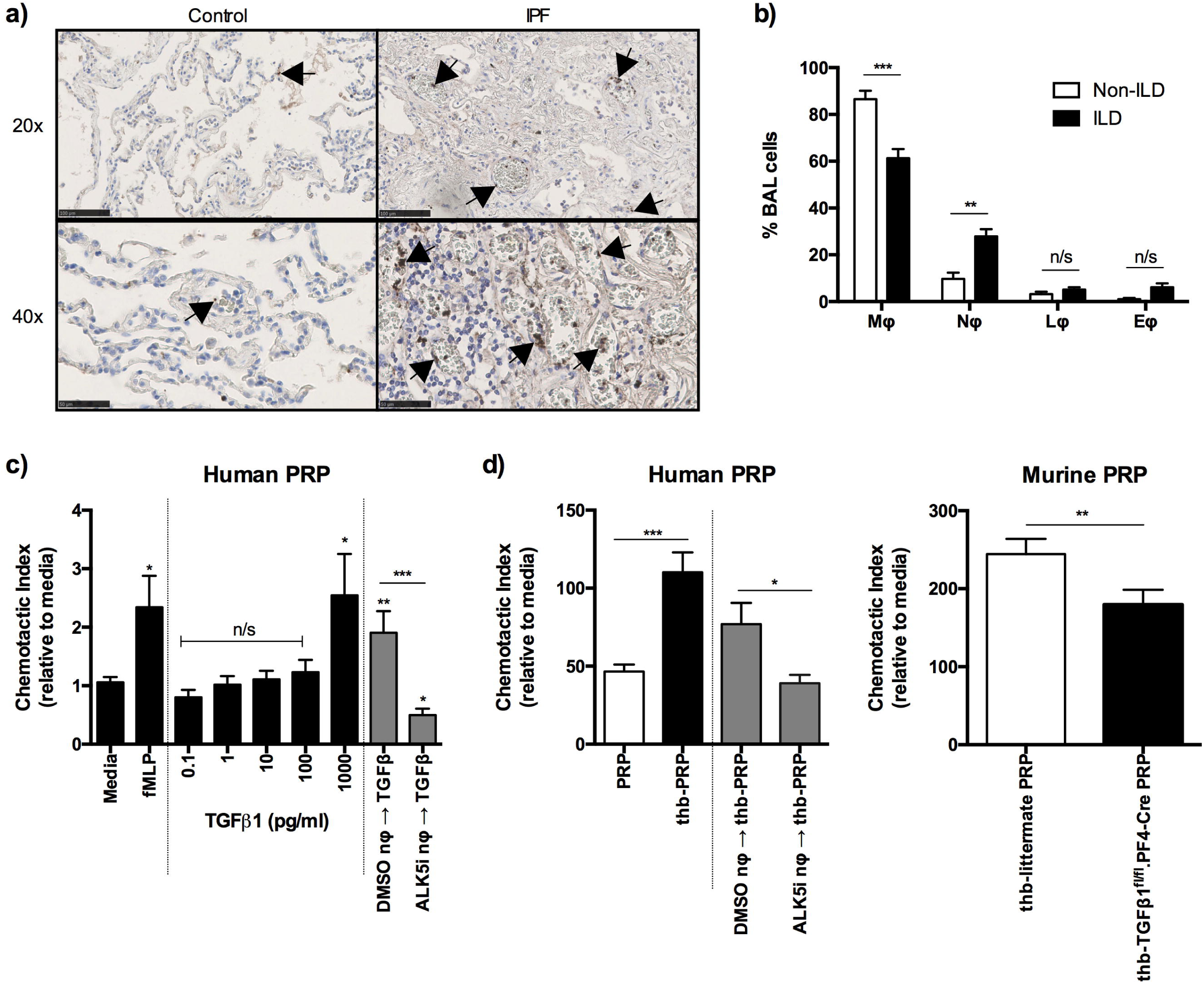
Platelet-derived TGFβ1 partially contributes to neutrophil chemotaxis. a) CD61^+^ platelets and platelet aggregates (indicated by the arrows) in control and IPF lung. Size denoted by scale bar (20x scale bar = 100μm, 40x scale bar = 50μm). b) BALF cell counts were determined in non-ILD (*n*=6) and ILD (*n*=12) patients by cytospin. c) Chemotaxis of untreated (black bars) or DMSO or ALK5 inhibitor (SB-525334) pre-treated (grey bars) human peripheral neutrophils (*n*=4 healthy donors) towards fMLP (100nM) or increasing concentrations of TGFβ1. d) Chemotaxis of untreated (white or black bars) or DMSO or ALK5 inhibitor (SB-525334) pre-treated (grey bars) human or murine peripheral neutrophils (*n*=4 healthy donors or *n*=4 littermate mice) towards unstimulated or thrombin (thb)-activated human or murine PRP. Statistical differences were determined using Mann-Whitney U test or 1-way ANOVA with Holm-Sidak post-hoc testing (n/s = not significant, \**p*<0.05, \*\**p*<0.01, \*\*\**p*<0.001).

We next verified whether platelet-derived TGFβ1 could induce human neutrophil migration. Significantly more neutrophils migrated towards thrombin-activated PRP than to unstimulated PRP (Fig. 4d). This effect was reduced when neutrophils were pre-treated with SB-525334 (Fig. 4d). This shows that human neutrophil chemotaxis towards thrombin-activated PRP is partly attributable to the presence of platelet-derived TGFβ1. This reduced migratory response was mirrored using TGFβ1^fl/fl^.PF4-Cre PRP with murine bone-marrow-derived neutrophils (Fig. 4d). These findings suggest that platelet-derived TGFβ1 partially contributes towards neutrophil migration *in vitro*.

### Platelet-derived TGFβ1 does not significantly induce neutrophil recruitment during pulmonary inflammation

To address whether platelet-derived TGFβ1 can influence cellular recruitment *in vivo*, littermate or TGFβ1^fl/fl^.PF4-Cre mice were treated with 50 IU bleomycin. During the peak of inflammation at day 6, littermate and TGFβ1^fl/fl^.PF4-Cre mice treated with bleomycin lost significantly more weight than saline treated mice (Fig. 5a). Increased neutrophils, macrophages and lymphocytes were in the BALF after bleomycin treatment compared to saline controls (Fig. 5b), with no significant difference between bleomycin-treated littermate or TGFβ1^fl/fl^.PF4-Cre mice (Fig. 5b). Flow cytometry was used to quantify neutrophil, alveolar and inflammatory macrophage recruitment into the lung (Fig. 5c). Bleomycin treatment led to elevated inflammatory macrophages recruitment compared to saline controls (Fig. 5c), with equivalent numbers between transgenic groups (Fig. 5c). To verify these results, we used another lung inflammation model using inhaled LPS. No significant differences in cellular recruitment into the lung tissue or BALF were found between littermate and TGFβ1^fl/fl^.PF4-Cre mice after 6h of LPS treatment (Supplementary Fig. 3). This suggests that despite platelet-derived TGFβ1 partially influencing neutrophil migration *in vitro*, it does not significantly drive cellular recruitment into the lung or airways *in vivo*.

**Figure 5.**
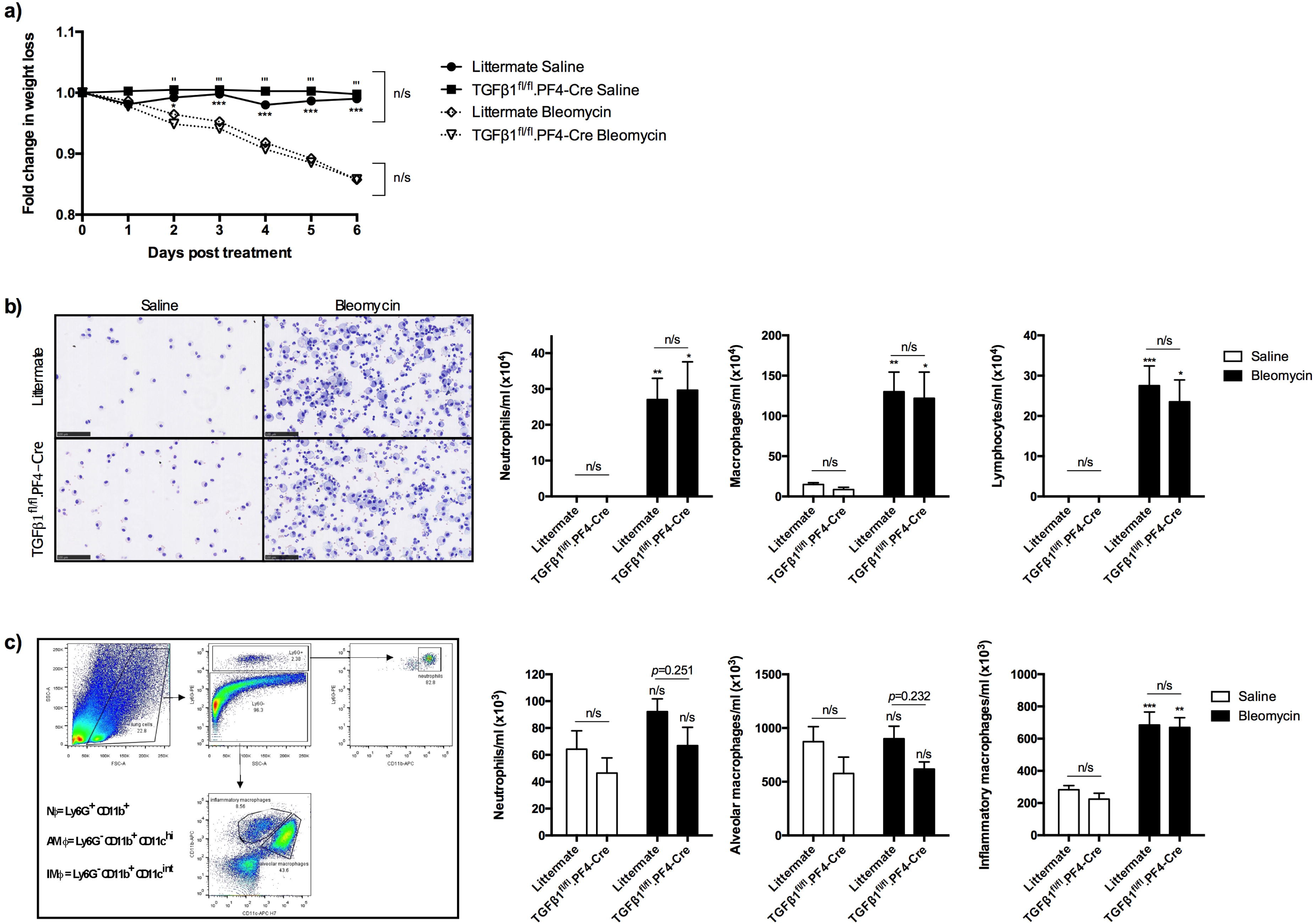
Platelet-derived TGFβ1 does not contribute to neutrophil recruitment during bleomycin-induced inflammation. a) TGFβ1^fl/fl^.PF4-Cre or littermates were given 50 IU bleomycin or saline (*n*=9 littermate saline, *n*=4 TGFβ1^fl/fl^.PF4-Cre saline, *n*=15 littermate bleomycin, *n*=8 TGFβ1^fl/fl^.PF4-Cre bleomycin) via an oropharyngeal route. Body weight was monitored for 6 days post instillation. b) Representative BALF cytospins after 6 days of treatment. Size denoted by scale bar (100μm). Cell populations in recovered BALF were quantified from cytospins. c) Representative flow cytometric gating strategy to distinguish recovered cell populations from lung homogenate. Cell populations in lung homogenate were quantified by flow cytometry. Statistical differences were determined using 1-way or 2-way ANOVA with Holm-Sidak post-hoc testing. On Fig. 5a, asterisks represent significance between treatments for littermate mice and apostrophes represent significance between treatments for TGFβ1^fl/fl^.PF4-Cre mice. On Fig. 5b and Fig. 5c, asterisks above the bars represent significance between saline or bleomycin group. Any other significant differences are indicated (n/s = not significant, \**p*<0.05, \*\**p*<0.01, \*\*\**p*<0.001).

### Platelet-derived mediators in ILD BALF may contribute to neutrophil recruitment and disease severity

As we previously observed an activated platelet signature in ILD blood, we next examined the BALF cellular composition in patients. ILD BALF contained significantly more platelets (Fig. 6a) and elevated total and active TGFβ1, PDGF-BB, CCL5 and MMP7 but not CXCL4 compared to non-ILD controls (Fig. 6b). ILD BALF MMP7 significantly positively correlated with paired CXCL4 levels (Fig. 6c), suggesting that more activated platelets may be linked with worse IPF disease severity. Total and active TGFβ1 and PDGF-BB did not correlate with CXCL4 (Fig. 6c), suggesting that activated platelets may not represent the key source of these pro-fibrotic mediators in the airways. We also observed strong positive correlation between CCL5, a neutrophil chemokine [28] with CXCL4 and MMP7 (Fig. 6d). Our data suggests that activated platelets may release CCL5 to drive the observed neutrophilic infiltration in ILD BAL.

**Figure 6.**
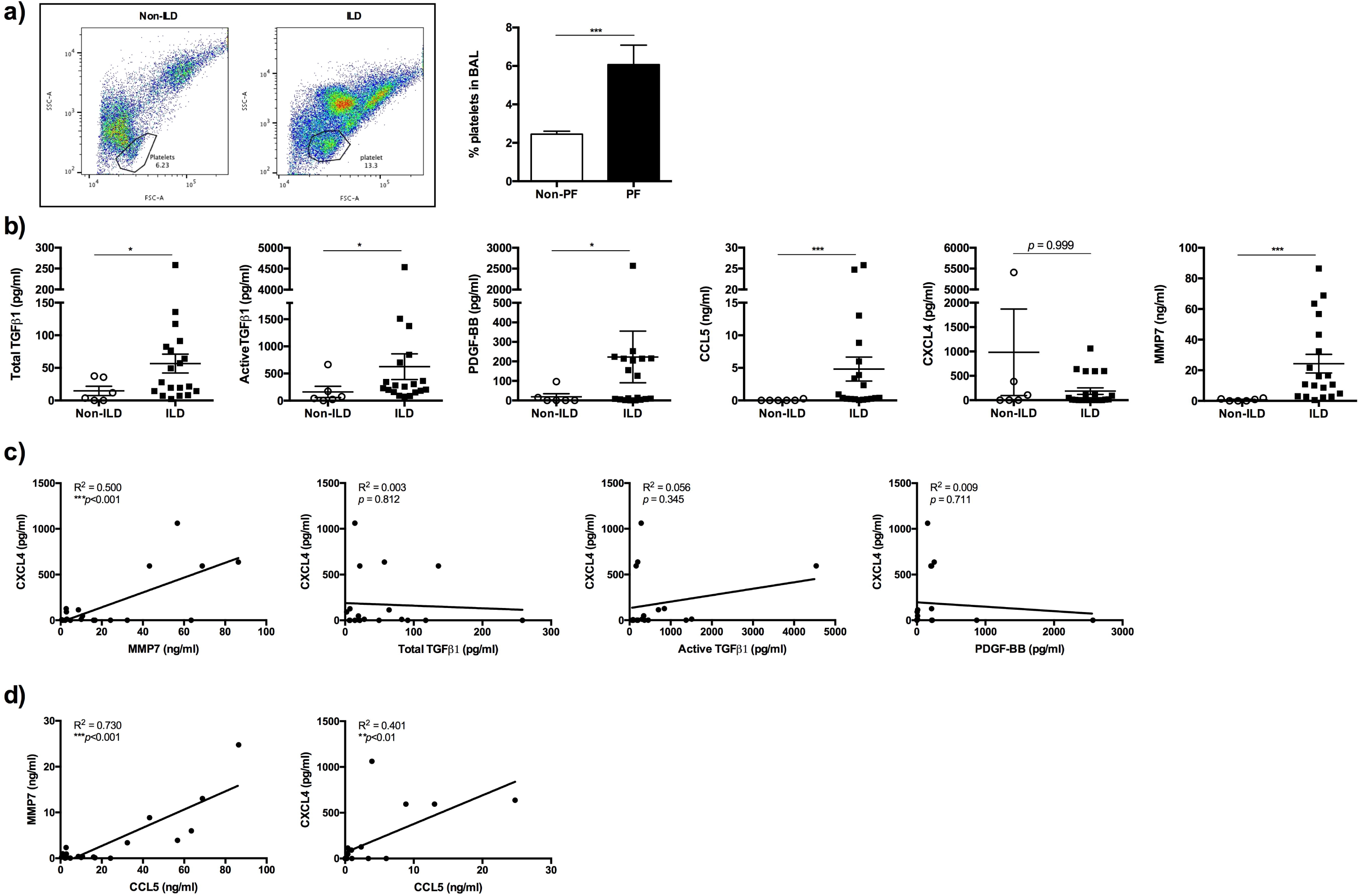
Platelet-derived mediators may contribute to neutrophil recruitment in IPF lung and disease severity. a) Representative flow cytometric plot showing platelet population in non-ILD and ILD BALF. Quantification of percentage of BALF platelets was done by flow cytometric analysis (*n*=3 non-ILD and *n*=6 ILD). b) Total or active TGFβ1, PDGF-BB, CCL5, CXCL4 and MMP7 concentrations were measured in non-ILD and ILD patients’ BALF (*n*=6 and 19 respectively) by ELISA or MLEC bioassay. c) Concentration of mediators in ILD patients’ BALF were correlated with paired CXCL4 (*n*=19). d) Concentration of CCL5 in ILD patients’ BALF were correlated with paired CXCL4 or MMP7 (*n*=19). Statistical differences were determined using Mann Whitney-U tests or linear regression and any significant differences are indicated (n/s = not significant, \**p*<0.05, \*\**p*<0.01, \*\*\**p*<0.001).

In summary, we found platelet-derived TGFβ1 has a redundant role in driving inflammation and fibrosis in bleomycin-induced PF. Our patient studies have revealed blood platelet counts predict survival in IPF and activated platelets in the lung may lead to neutrophil recruitment and drive human PF disease progression through the release of mediators such as CCL5 and not TGFβ1.

## Discussion

Increased platelet reactivity and trapping within the lung have been previously described in IPF clinical and animal studies [15, 16]. Whilst, these reports were unable to establish a direct causative relationship between platelets or platelet-derived products with IPF disease progression, the aim of this study was to determine the contribution of platelet-derived TGFβ1 in mediating lung inflammation and fibrosis *in vivo*.

Our findings of elevated MMP7 in ILD/IPF plasma compared to non-ILD controls agrees with literature proposing MMP7 as a IPF plasma biomarker [26]. Our patient cohort also displayed an activated blood platelet signature correlating with increased total and active TGFβ1, suggesting platelets as a likely source of this pro-fibrotic cytokine in IPF. Interestingly, we did not find any significant difference in the blood platelet counts between our patient groups (data not shown), whereas Ntolios *et al*., have previously showed IPF patients have lower blood platelet counts compared to controls [14]. This discrepancy could be due to sample size and choice of patients recruited.

Recent evidence proposes the lung as a major site for platelet biogenesis in mice with a distinct megakaryocyte subset, residing within the lung interstitium [29]. It is unknown whether megakaryopoiesis or the presence of interstitial megakaryocytes also occur in human lungs, although megakaryocytes have been found in human lung capillaries [30]. We readily detected platelets in the vasculature of lung and BALF of ILD patients, however, we did not find any megakaryocytes within the blood vessels or interstitium of control or IPF lung.

Platelets were detected in bleomycin-treated littermate and TGFβ1^fl/fl^.PF4-Cre lungs, however the degree of fibrosis was equivalent in both treatment groups, suggesting that platelet-derived TGFβ1 is redundant in this animal PF model. PRP can enhance fibroblast cell growth and contraction of collagen matrix [31, 32], although the relative contribution of platelet-derived TGFβ1 was not investigated. Activated platelets can secrete over 300 proteins [33] and PRP contains key pro-fibrotic growth factors such as PDGF [34]. Therefore, it is not inconceivable that these growth factors may play a dominant pro-fibrotic role in the absence of platelet-derived TGFβ1 in TGFβ1^fl/fl^.PF4-Cre mice. Studies inhibiting PDGF-BB signalling with Imatinib (a tyrosine kinase inhibitor of PDGF-R) attenuated bleomycin-induced PF [35], emphasising the importance of studying other platelet-derived products in mitigating PF pathology.

We have shown PRP is a potent neutrophil chemoattractant and platelet-derived TGFβ1 partially contributed to neutrophil migration *in vitro*. TGFβ1 has previously been reported as a neutrophil chemoattractant [36], but to our best knowledge, this is the first instance that platelet-derived TGFβ1 has been linked to neutrophil chemotaxis. We observed no difference in the number or percentage of neutrophils recruited into the lung or BALF of littermate or TGFβ1^fl/fl^.PF4-Cre mice after bleomycin or LPS challenge. This suggests that platelet-derived TGFβ1 has a redundant role in neutrophil migration *in vivo*. CXCL5 and CXCL7 are important platelet-derived neutrophil chemoattractants in a murine metastasis model [37], and it is probable that these and other platelet-derived neutrophil chemoattractants, such as CCL5 [38] could compensate for the absence of TGFβ1 in TGFβ1^fl/fl^.PF4-Cre mice to recruit neutrophils as suggested by our clinical observations.

Of importance, it should be noted that whilst, bleomycin remains the gold standard animal model for PF research, it does not fully recapitulate the human disease. The degree of lung fibrosis in mice is dependent on bleomycin dose [39], whilst IPF is a non-resolving progressive disease. Our patient analysis showed a novel significant correlation between mortality and elevated platelet counts, along with an activated platelet signature associated with increased TGFβ1 in ILD blood to strongly suggest a detrimental role for platelets in ILD. Further investigation of these abnormal platelet responses represents an exciting new avenue of IPF research. In particular, further validation of our new observations in another patient cohort could lead to the development of platelet count or activation as a prognostic or treatment response biomarker in IPF and even as a potential treatment target.

Overall, the novel findings we present here offer a better understanding of platelets and platelet-derived TGFβ1 in both human IPF disease and animal models and further studies may facilitate a more focused cell-targeted approach to TGFβ1 inhibition.

## Acknowledgements

We thank Prof. Richard Hynes (MIT, USA) for providing the TGFβ1^fl/fl^.PF4-Cre mice for our study.

## Financial Support

This work was undertaken at UCLH/UCL, which received a proportion of the funding from the UK’s Department of Health’s NIHR Biomedical Research Centre’s fund scheme. This work was also supported by the Medical Research Council (MRC) (grant codes G0800340 and MR/K004158/1) and a European Commission FP7 award PIEF GA-2012-326928 to Dr Rebeyrol. The authors declare no competing financial interest.

## References

1. Mikolasch TA, Garthwaite HS, Porter JC. Update in diagnosis and management of interstitial lung disease. Clin Med 2016: 16(6): s71–78.

2. Gross TJ, Hunninghake GW. Idiopathic Pulmonary Fibrosis. N Engl J Med 2001: 345(7): 517–525.

3. Biernacka A, Dobaczewski M, Frangogiannis NG. TGF-β signaling in fibrosis. Growth Factors 2011: 29(5): 196–202.

4. Khalil N, Parekh TV, O’Connor RO, Antman N, Kepron W, Yehaulaeshet T, Xu YD, Gold LI. Regulation of the effects of TGF-β1 by activation of latent TGF-β1 and differential expression of TGF-β receptors (TβR-I and TβR-II) in idiopathic pulmonary fibrosis. Thorax 2001: 56: 907–915.

5. Santana A, Saxena B, Noble NA, Gold LI, Marshall BC. Increased Expression of Transforming Growth Factor β Isoforms (β1, β2, β3) in Bleomycin-induced Pulmonary Fibrosis. Am J Respir Cell Mol Biol 1995: 13: 34–44.

6. Xaubet A, Marin-Arguedas A, Lario S, Ancochea J, Morell F, Ruiz-Manzano J, Rodriguez-Becerra E, Rodriguez-Arias JM, Inigo P, Sanz S, Campistol JM, Mullol J, Picado C. Transforming growth factor-beta1 gene polymorphisms are associated with disease progression in idiopathic pulmonary fibrosis. Am J Respir Crit Care Med 2003: 168(4): 431–435.

7. Khalil N, Bereznay O, Sporn M, Greenberg AH. Macrophage production of transforming growth factor β and fibroblast collagen synthesis in chronic pulmonary inflammation. J Exp Med 1989: 170(3): 727–737.

8. Zhang K, Flanders KC, Phan SH. Cellular Localization of Transforming Growth Factor-β Expression In Bleomycin-Induced Pulmonary Fibrosis. Am J Pathol 1995: 147(2): 352–361.

9. Celada LJ, Kropski JA, Herazo-Maya JD, Luo W, Creecy A, Abad AT, Chioma OS, Lee G, Hassell NE, Shaginurova GI, Wang Y, Johnson JE, Kerrigan A, Mason WR, Baughman RP, Ayers GD, Bernard GR, Culver DA, Montgomery CG, Maher TM, Molyneaux PL, Noth I, Mutsaers SE, Prele CM, Peebles Jr RS, Newcomb DC, Kaminski N, Blackwell TS, Van Kaer L, Drake WT. PD-1 up-regulation on CD4+ T cells promotes pulmonary fibrosis through STAT3-mediated IL-17A and TGF-β1 production. Sci Transl Med 2018: 10: eaar8356.

10. Assoian RK, Komoriya A, Meyers CA, Miller DM, Sporn MB. Transforming Growth Factor-β in Human Platelets. Identification of a major storage site, purification, and characterization. J Biol Chem 1983: 258(11): 155–160.

11. Fox KA, Kirwan DE, Whittington AM, Krishnan N, Robertson BD, Gilman RH, Löpez JW, Singh S, Porter JC, Friedland JS. Platelets Regulate Pulmonary Inflammation and Tissue Destruction in Tuberculosis. Am J Respir Crit Care Med 2018: 198(2): 245–255.

12. Watanabe M, Murata S, Hashimoto I, Nakano Y, Ikeda O, Aoyagi Y, Matsuo R, Fukunaga K, Yasue H, Ohkohchi N. Platelets contribute to the reduction of liver fibrosis in mice. J Gastroenterol Hepatol 2009: 24: 78–89.

13. Meyer A, Wang W, Qu J, Croft L, Degen JL, Coller BS, Ahamed J. Platelet TGF-β1 contributions to plasma TGF-β1, cardiac fibrosis, and systolic dysfunction in a mouse model of pressure overload. Blood 2012: 119(4): 1064–1074.

14. Ntolios P, Papanas N, Nena E, Boglou P, Koulelidis A, Tzouvelekis A, Xanthoudaki M, Tsigalou C, Froudarakis ME, Bouros D, Mikhailidis DP, Steiropoulos P. Mean Platelet Volume as a Surrogate Marker for Platelet Activation in Patients With Idiopathic Pulmonary Fibrosis. Clin Appl Throm Haem 2016: 22(4): 346–350.

15. Crooks MG, Fahim A, Naseem KM, Morice AH, Hart SP. Increased platelet reactivity in idiopathic pulmonary fibrosis is mediated by a plasma factor. PLoS One 2014: 9(10): e111347.

16. Piguet PF, Vesin C. Pulmonary platelet trapping induced by bleomycin: correlation with fibrosis and involvement of the beta 2 integrins. Int J Exp Path 1994: 75: 321–328.

17. Sreeramkumar V, Adrover JM, Ballesteros I, Cuartero MI, Rossaint J, Bilbao I, Nácher M, Pitaval C, Radovanovic I, Fukui Y, McEver RP, Filippi MD, Lizasoain I, Ruiz-Cabello J, Zarbock A, Moro MA, Hidalgo A. Neutrophils scan for activated platelets to initiate inflammation. Science 2014: 346(6214): 1234–1238.

18. Kinder BW, Brown KK, Schwarz MI, Ix JH, Kervitsky A, King TE, Jr. Baseline BAL neutrophilia predicts early mortality in idiopathic pulmonary fibrosis. Chest 2008: 133(1): 226–232.

19. Chua F, Dunsmore SE, Clingen PH, Mutsaers SE, Shapiro SD, Segal AW, Roes J, Laurent GJ. Mice lacking neutrophil elastase are resistant to bleomycin-induced pulmonary fibrosis. Am J Pathol 2007: 170(1): 65–74.

20. Chrysanthopoulou A, Mitroulis I, Apostolidou E, Arelaki S, Mikroulis D, Konstantinidis T, Sivridis E, Koffa M, Giatromanolaki A, Boumpas DT, Ritis K, Kambas K. Neutrophil extracellular traps promote differentiation and function of fibroblasts. J Pathol 2014: 233(3): 294–307.

21. Labelle M, Begum S, Hynes RO. Direct signaling between platelets and cancer cells induces an epithelial-mesenchymal-like transition and promotes metastasis. Cancer Cell 2011: 20(5): 576–590.

22. Abe M, Harpel JG, Metz CN, Nunes I, Loskutoff DJ, Rifkin DB. An Assay for Transforming Growth Factor-β Using Cells Transfected with a Plasminogen Activator Inhibitor-1 Promoter-Luciferase Construct. Anal Biochem 1994: 216: 276–284.

23. Scotton CJ, Hayes B, Alexander R, Datta A, Forty EJ, Mercer PF, Blanchard A, Chambers RC. Ex vivo micro-computed tomography analysis of bleomycin-induced lung fibrosis for preclinical drug evaluation. Eur Respir J 2013: 42(6): 1633–1645.

24. Campa JS, McAnulty RJ, Laurent GJ. Application of High-Pressure Liquid Chromatography to Studies of Collagen Production by Isolated Cells in Culture. Anal Biochem 1990: 186: 257–263.

25. Rucinski B, Niewiarowski S, Strzyzewski M, Holt JC, Mayo KH. Human platelet factor 4 and its C-terminal peptides: heparin binding and clearance from the circulation. Thromb Haemost 1980: 63(3): 493–498.

26. Rosas IO, Richards TJ, Konishi K, Zhang Y, Gibson KF, Lokshin AE, Lindell KO, Cisneros J, MacDonald SD, Pardo A, Sciurba F, Dauber J, Selman M, Gochuico BR, Kaminski N. MMP1 and MMP7 as Potential Peripheral Blood Biomarkers in Idiopathic Pulmonary Fibrosis. Plos Med 2008: 5(4): e93.

27. Grainger DJ, Wakefield L, Bethell HW, Farndale RW, Metcalfe JC. Release and activation of platelet latent TGF-β in blood clots during dissolution with plasmin. Nat Med 1995: 1(9): 932–937.

28. Pan Z, Parkyn L, Ray A, Ray P. Inducible lung-specific expression of RANTES: preferential recruitment of neutrophils. Am J Physio Lung Cell Mol Physiol 2000: 276: L658–L666.

29. Lefrançais E, Oritx-Muñoz G, Caudrillier A, Mallavia B, Liu F, Sayah DM, Thornton EE, Headley MB, David T, Coughlin SR, Krummel MF, Leavitt AD, Passegué E, Looney MR. The lung is a site of platelet biogenesis and a reservoir for haematopoietic progenitors. Nature 2017.

30. Ouzegdouh Y, Capron C, Bauer T, Puymirat E, Diehl J, Martin JF, Cramer-Bordé E. The physical and cellular conditions of the human pulmonary circulation enable thrombopoesis. Exp Hematol 2018.

31. Passaretti F, Tia M, D’Esposito V, De Pascale M, Del Corso M, Sepulveres R, Liguoro D, Valentino R, Beguinot F, Formisano P, Sammartino G. Growth-promoting action and growth factor release by different platelet derivatives. Platelets 2014: 25(4): 252–256.

32. Zagai U, Fredriksson K, Rennard SI, Lundahl J, Sköld CM. Platelets stimulate fibroblast-mediated contraction of collagen gels. Respir Res 2003: 4: 13.

33. Coppinger JA, Cagney G, Toomey S, Kislinger T, Belton O, McRedmond JP, Cahill DJ, Emili A, Fitzgerald DJ, Maguire PR. Characterization of the proteins released from activated platelets leads to localization of novel platelet proteins in human atherosclerotic lesions. Blood 2004: 103: 2096–2104.

34. Antoniades HN, Scher CD, Stiles CD. Purification of human platelet-derived growth factor. Proc Natl Acad Sci U S A 1979: 76(4): 1809–1813.

35. Aono Y, Nishioka Y, Inayama M, Ugai M, Kishi J, Uehara H, Izumi K, Sone S. Imatinib as a Novel Antifibrotic Agent in Bleomycin-induced Pulmonary Fibrosis in Mice. Am J Respir Crit Care Med 2005: 171: 1279–1285.

36. Reibman J, Meixler S, Lee TC, Gold LI, Cronstein BN, Haines KA, Kolasinski SL, Weissmann G. Transforming growth factor β1, a potent chemoattractant for human neutrophils, bypasses classic signal-transduction pathways. Proc Natl Acad Sci U S A 1991: 88: 6805–6809.

37. Labelle M, Begum S, Hynes RO. Platelets guide the formation of early metastatic niches. Proc Natl Acad Sci U S A 2014: 111(30): E3053–3061.

38. Hwaiz R, Rahman M, Syk I, Zhang E, Thorlacius H. Rac1-dependent secretion of platelet-derived CCL5 regulates neutrophil recruitment via activation of alveolar macrophages in septic lung injury. J Leukoc Biol 2015.

39. Williamson JD, Sadofsky LR, Hart SP. The pathogenesis of bleomycin-induced lung injury in animals and its applicability to human idiopathic pulmonary fibrosis. Exp Lung Res 2015: 41(2): 57–73.

