## Supplemental Methods and Figures for "Platelet counts predict prognosis in IPF, but are not the main source of pulmonary TGFβ1"

**Supplementary Methods**

**Patients recruitment for survival analysis**

For the survival analysis, unselected, consecutive IPF patients from the UCLH ILD Service were followed prospectively for mortality based on blood platelet counts. The diagnosis of IPF was made following clinico-radiological multidisciplinary team review. Survival time was taken from point of first presentation to UCLH to time of death or lung transplant. Patients with platelet counts outside the normal range, known haematological disorders or treatments known to affect platelet count were excluded. Patients were divided into 3 categories with cut-offs determined bias free as the lower, middle and upper thirds. The cut-offs were 207x10^9^/L and 267x10^9^/L respectively.

**Patient BALF & plasma collection & processing**

Paired blood and BALF samples were collected from patients consented for a routine bronchoscopy for diagnostic reasons. Lavages were collected bronchoscopically after saline instillation, and aliquots of BALF were frozen after centrifugation for later cytokine analysis. The remaining BAL cell pellet was resuspended in PBS before being cytocentrifuged and differentially stained with a Rapid Romanowsky Stain Kit (TCS Biosciences Ltd, Botolph Claydon, UK). Cell counts were performed using a conventional brightfield microscope. Whole blood was collected in EDTA vacutainers before centrifugation to collect the plasma for later cytokine analysis. Percentage of platelets in BALF was quantified on a BD FACSVerse flow cytometer (BD Biosciences, San Jose, USA) and analysed with FlowJo Version 10 software (FlowJo LLC, Ashland, USA).

**Neutrophil chemotaxis assay**

Human or murine PRP were obtained by collecting whole blood in EDTA, followed by centrifugation for 8 min at 800 *g*. PRP was activated with 0.5 U/ml thrombin (Sigma-Aldrich, Gillingham, UK) for 20 min at RT prior to centrifugation at 2800 *g* for 7 min to remove the platelet pellet. Murine bone marrow-derived neutrophils were collected by flushing the femurs of transgenic mice with 3 washes of 1 ml PBS. Human neutrophils (*n*=4 healthy donors) were obtained from citrated peripheral blood collected from healthy donors after an initial 6% dextran (Sigma-Aldrich) sedimentation step to remove RBC. Murine bone marrow-derived neutrophils (*n*=4 littermate controls) or human neutrophils were isolated from a Percoll Plus (Sigma-Aldrich) gradient by centrifugation at 700 *g* for 30 min before labelling with 1 μM CMFDA (Invitrogen, Carlsbad, USA) in HBSS- for 30 min at 37°C. Cells were washed once in HBSS- and resuspended at 5x10^6^ cells/ml and 2.5x10^6^ cells/ml in HBSS+ for human or murine neutrophils respectively. 100 μl neutrophils were added to the top of 8 μm PET transwells insert (Corning, New York, USA) and chemoattractants: 100 nM fMLP (Sigma-Aldrich), TGFβ1 (R&D systems), 10 ng/ml LTB_4_ (Cambridge Biosciences, Bar Hill, UK) or pre-diluted PRP (*n*=4) added to the top or bottom of the insert. In ALK5 inhibition studies, neutrophils were pre-treated with 1 μM SB-525334 (Sigma-Aldrich) for 30 min before migration. Neutrophils were allowed to migrate for 1 hour at 37°C before collection and read on a BD FACSVerse flow cytometer and analysed with FlowJo software. The degree of migration was expressed as the Chemotactic Index (number of cells migrating in response to the stimulus ÷ number of cells migrating in response to media control).

**Histology & IHC**

Murine or human lung samples were processed for histological analysis as described (Smoktunowicz *et al*., 2015. Dis Model Mech; 9:1129-1139). In brief, re-hydrated micro-CT processed murine lungs or human lung biopsies were embedded in paraffin wax blocks. 5μm sections were stained with Hematoxyline & Eosin (H&E) or Martius Scarlet Blue (MSB) using an automated slide stainer (Sakura Tissue-Tek DRS 2000, Alphen aan den Rijn, The Netherlands). IHC was performed for the presence of murine CD61 (AB-7773, Sigma-Aldrich) or human CD61 (2f2, Leica Biosystems, Wetzlar, Germany) under heat-induced epitope retrieval methods with citrate buffer pH 6.0. Staining was developed using HRP-conjugated secondary antibody and NovaRED Peroxide (HRP) substrate kit (all from Vector Laboratories, Burlingame, USA) for murine tissue or DAB for human tissue (Leica Biosystems) and sections counter-stained with Haematoxylin. All sections were scanned on a Nanozoomer Digital Slide Scanner and images analysed using NDP.view software (both from Hamamatsu Corporation, Hamamatsu City, Japan).

**LPS animal model of inflammation**

15-20 week old male and female TGFβ1^fl/fl^.PF4-Cre or littermate control mice were given 3.75 μg/mouse LPS (Sigmal-Aldrich) in a volume of 50 μl saline or saline only via an intranasal route. Mice were monitored for disease severity and culled 6 hours post instillation. BAL and lungs samples were collected for cytospins, cytokine analysis and flow cytometry as described above.

**Supplementary Table 1: Summary of IPF patient cohort for prospective mortality study.**

| **Platelet Strata** | **Mean Age (SD)** | **Mean FVC^*^ % predicted (SD)** | **% Male** |
| --- | --- | --- | --- |
| Group 1 (*n*=71) | 73.3 (10.3) | 77.9 (21.8) | 77.5% |
| Group 2 (*n*=72) | 73.9 (8.4) | 75.9 (18.6) | 86.1% |
| Group 3 (*n*= 71) | 72.8 (8.3) | 73.4 (19.8) | 74.7% |

* FVC Forced Vital Capacity

**Supplementary table 2. Summary of Non-ILD and ILD patient cohort for cytokine analysis.**

| **Characteristics** | **Non-ILD cohort** | **ILD cohort** |
| --- | --- | --- |
| Blood samples recruited | 9 | 47 |
| Mean Age (SD) | 59.8 (15.9) | 71.2 (8.1) |
| % Male | 55.6% | 68.1% |
| Diagnosis | Haemoptysis with no lung pathology (*n*=5),  Healthy control (*n*=2),  Previous schwannoma (*n*=1),  Asthma/COPD (*n*=1) | IPF (*n*=24),  Chronic hypersensitivity pneumonitis (HP) (*n*=5),  Unclassifiable fibrotic ILD (*n*=3),  Rheumatoid arthritis (RA)-ILD (*n*=3),  Chronic HP/Sarcoid (*n*=2),  Nonspecific interstitial pneumonia (NSIP) (*n*=6),  Inflammatory drug induced ILD (*n*=1),  Fibrosing organising pneumonia (FOP) (*n*=1),  Usual interstitial pneumonia (UIP) (*n*=1),  Combined PF & emphysema (CPFE) (*n*=1) |
| Lavage samples recruited | 6 | 19 |
| Mean Age (SD) | 51.0 (11.0) | 68.3 (5.7) |
| % Male | 50.0% | 73.7% |
| Diagnosis | Haemoptysis with no lung pathology (*n*=5),  Previous schwannoma (*n*=1) | IPF (*n*=6),  Unclassifiable fibrotic ILD (*n*=4),  Chronic HP/Sarcoid (*n*=2),  NSIP (*n*=3),  UIP (*n*=2),  Inflammatory drug induced ILD (*n*=1),  Chronic HP (*n*=1) |

**Supplementary figure 1. Activated human platelets are a source of active TGFβ1**

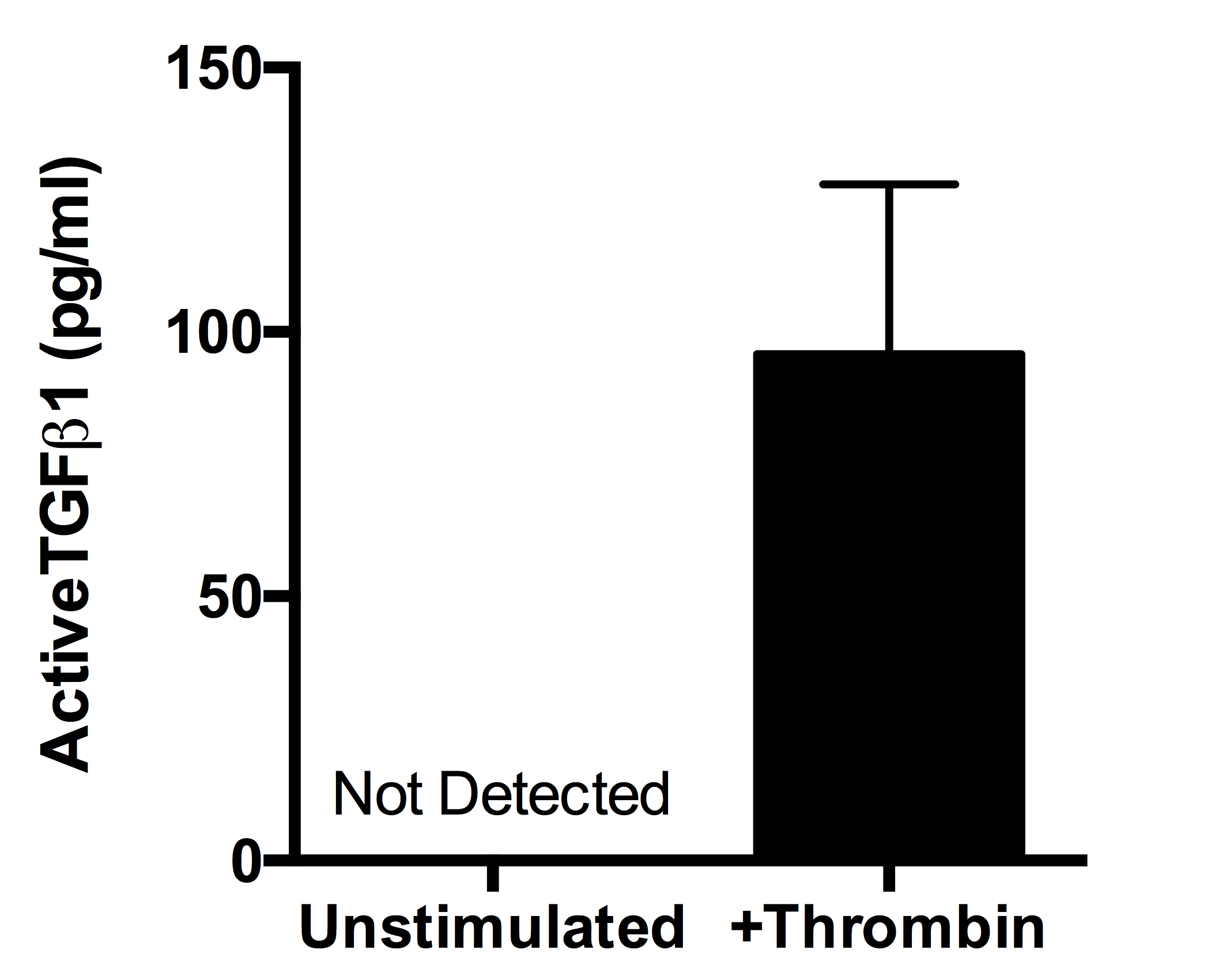

Quantification of active TGFβ1 by MLEC bioassay in unstimulated or thrombin-treated human PRP (*n*=3 healthy donors).

**Supplementary figure 2. Littermate and TGFβ1^fl/fl^.PF4.Cre mice exhibit similar degrees of inflammation and fibrosis after *in vivo* bleomycin challenge*.***

TGFβ1^fl/fl^.PF4-Cre (*n*=10) or littermate control mice (*n*=8) were given 25 IU bleomycin via an oropharyngeal route. Lung tissue was harvested 28 days later for histological analysis. Littermate (left column) or TGFβ1^fl/fl^.PF4-Cre lungs (right column) were processed for H&E, modified trichrome (MT; blue=collagen, yellow=RBC, red=cytoplasm, dark red=nuclei), CD61 or secondary antibody (2’Ab) only control IHC staining (positive staining = brown, arrows denote platelets and platelet aggregates). Images are shown at 20x objective as denoted by the scale bar.

**Supplemental Figure 3. Platelet-derived TGFβ1 does not play a significant role in mediating inflammation after *in vivo* LPS challenge**

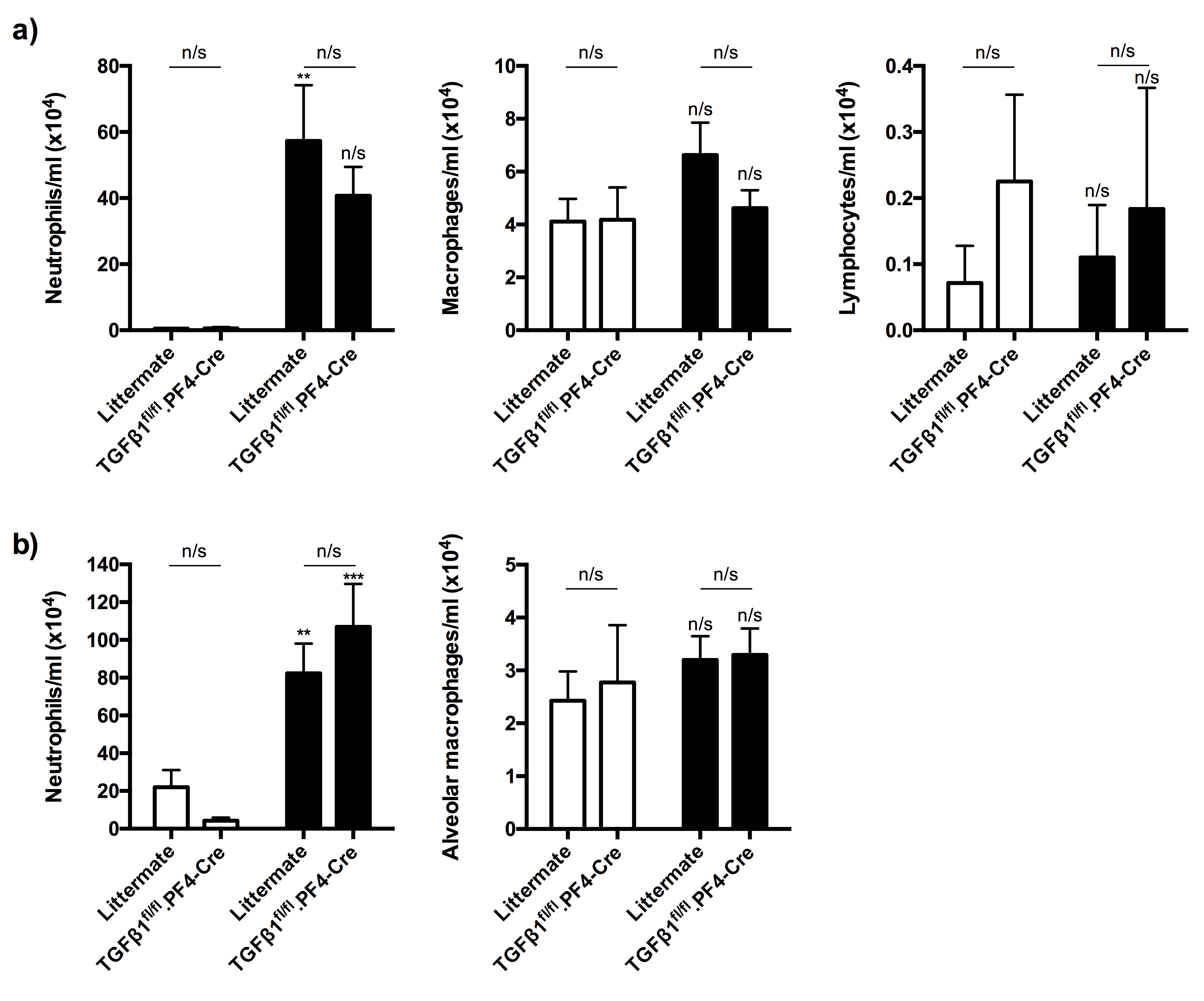

TGFβ1^fl/fl^.PF4-Cre or littermate control mice were given saline (*n*=7 littermate or *n*=4 TGFβ1^fl/fl^.PF4-Cre) or 3.75 μg/mouse LPS (*n*=10 littermate or *n*=6 TGFβ1^fl/fl^.PF4-Cre) via intranasal administration. BALF and lung tissue were harvested 6 hours later to determine the degree of inflammation.

a) Different subsets of cells were counted by cytospin from the recovered BALF.

b) Different subsets of cells were counted by flow cytometry from the recovered lung homogenate.

Any statistical differences were determined using a 2-way ANOVA test with Holm-Sidak post-hoc testing. Asterisks above the bars represent significance between saline & LPS treatment (n/s = not significant, ***p*<0.01, ****p*<0.001).
